# Transcriptome-based cell type assignment for kidney cell culture models

**DOI:** 10.64898/2026.03.30.715265

**Authors:** Mona Schoberth, Samuel Böhm, Oleg Borisov, Yong Li, Gabriele Greve, Bayram Edemir, Owen M. Woodward, Hyun Jun Jung, Frank Hutter, Lukas Westermann, Anna Köttgen, Pascal Schlosser, Michael Köttgen, Stefan Haug

## Abstract

**Background:** Kidney cell lines are widely used to model kidney physiology and disease; however, their gene expression profiles may differ from primary cells due to immortalization, culture conditions, or experimental treatments. Determining whether a cell line resembles its native cell type is critical for interpreting in vitro findings. We developed a transcriptome-based approach that matches bulk RNA-seq data from kidney cell lines, primary cells, or tissues to reference cell types derived from single-cell RNA-seq (scRNA-seq) datasets.

**Methods:** Reference transcriptomic profiles were generated from two human and two murine kidney scRNA-seq datasets by pseudobulk aggregation. Bulk RNA-seq data from microdissected kidney tissue, non-kidney negative controls, and kidney cell lines were matched to these references using three statistical similarity measures (Spearman correlation, Euclidean distance, Poisson distance) and three machine learning classifiers (Random Forest, XGBoost, TabPFN). Each was assessed with global gene expression, curated kidney marker gene lists, and the most variable genes. Matching accuracy was evaluated through a three-step validation strategy: within-dataset matching, cross-reference comparison, and validation against primary kidney tissue and negative controls.

**Results:** Gene expression rank-based Spearman correlation and TabPFN, a foundation model for tabular data, emerged as the most accurate and specific approaches, particularly with curated kidney marker gene lists. Both methods correctly identified microdissected kidney tubule segments and were robust against non-kidney negative controls. Applied to commonly used kidney cell lines, OK cells retained proximal tubule identity, particularly under shear stress, while other proximal tubule lines (HK-2, HKC-8, HKC-11) showed inconsistent matching. Collecting duct-derived mIMCD-3 maintained stable similarity across passages, culture conditions, and genetic modifications.

**Conclusion:** We provide two complementary implementations: CellMatchR, an accessible web-based tool using Spearman correlation for routine use, and comprehensive scripts for TabPFN-based matching (link will be added after peer reviewed publication). Together, these resources enable researchers to make informed decisions about kidney cell culture model selection, interpretation, and stability.

**Translational Statement:** Kidney cell lines are fundamental tools in nephrology research, yet their transcriptomic similarity to native cell types is rarely validated systematically. We demonstrate that combining bulk RNA-seq data with single-cell reference datasets enables robust assessment of cell line identity using gene expression-rank-based correlation and machine learning approaches. By providing a comprehensive evaluation of matching methods, curated kidney marker gene lists, and reference datasets, our study serves as both a practical resource and a methodological framework for the kidney research community, facilitating informed selection of cell culture models, quality control of experimental conditions, developing new experimental cell culture models, and more reliable translation of *in vitro* findings to kidney physiology and disease.

## Introduction

Cell culture models are widely used in kidney research to study physiological and pathophysiological processes relevant to diseases such as acute^1,2^ and chronic^3^ kidney injury, cystic diseases^4^ and cancer^5,6^. At least 16 tubular cell types with distinct functions and transcriptomic profiles exist^7^, and immortalized lines are most commonly derived from proximal tubule or collecting duct^8^. However, establishing such cultures may result in de-differentiation, and loss of transporters and cell polarization^8^. Cell morphology and gene expression are further influenced by passage number^9^, culture conditions and the source of the cell line^8^. One of the most widely used proximal tubule cell lines, HK-2, was derived from a healthy adult kidney^10^ but has lost expression of key proximal tubule transporters (OAT1, OAT3, ABCG1)^11^, limiting its value as a physiological model. More broadly, none of the 14 proximal tubule lines examined in a publication by Khundmiri *et al*. retained strong similarity to native cells based on the presence of cell type-specific transcripts^8^.

To determine which reference cell type a cell line most closely resembles, current approaches typically rely on individual marker genes, which may miss broader transcriptomic changes affecting cellular phenotype. Tools such as CellNet^12^ evaluate engineered cells against tissue types using gene regulatory networks, and single-cell label transfer methods (e.g., SingleR^13^) enable automated annotation between scRNA-seq datasets. Deconvolution methods^14,15^ estimate cell type proportions in heterogeneous bulk samples but assume cell type mixtures which is typically not applicable to homogeneous cell lines. No framework exists for matching bulk RNA-seq from cell lines to kidney-specific cell types using large sets of genes. Given that RNA sequencing (RNA-seq) data are routinely generated and multiple human and mouse kidney single-cell RNA-seq (scRNA-seq) references are now available^16–19^, we developed a transcriptome-based approach to match cell lines to kidney reference cell types derived from single-cell datasets. We systematically evaluated different similarity measures, machine learning approaches, gene sets, and scRNA-seq reference datasets for their accuracy in assigning primary kidney cells and cell lines to their cell type with the best transcriptomic match. We provide comprehensive online resources implementing the most reliable approaches (https://github.com/genepi-frei-burg/CellMatchR), enabling researchers to assess the cellular identity of kidney-specific in vitro models without extensive bioinformatic knowledge.

## Methods

Reference transcriptomic profiles for kidney cell types were derived from two murine (Ransick *et al*.^16^, Park *et al*.^17^) and two human scRNA-seq datasets (KPMP^19^ and Zhang *et al*.^18^) (**Supplementary Table 1**). For each dataset, we computed cell type-specific reference transcriptomes by summing the counts for each gene from all cells assigned to a given cell type and then normalizing to counts per million (CPM) to account for differences in cell numbers and sequencing depth.

We analyzed multiple publicly available and in-house bulk RNA-seq datasets from kidney tissues and cell lines (**Supplementary Table 2)**, which were matched to the kidney cell type references using four gene sets: all shared genes (global expression), curated markers for all kidney cells (mg_all, n=584) and kidney tubule-specific cells (mg_tub, n=310, **Supplementary Table 3**), and the 1000 most variable genes per reference. Based on each gene set, cell type similarity was assessed using Spearman rank correlation, Euclidean distance, or Poisson distance^20^ (“similarity methods”), with and without dimensional reduction by PCA or UMAP (**Supplementary Methods**). Additionally, three machine learning (ML) classifiers (Random Forest^21^, XGBoost^22^, and TabPFN v2.5^23,24^) were evaluated using identical gene sets for comparability with the exception of TabPFN, which was not tested using all genes as the model capacity is limited to a maximum of 2000 features. ML methods were also tested with pooled reference data to leverage training across multiple datasets. Matching accuracy was defined as the percentage of samples where the biologically correct reference cell type achieved the highest similarity score.

To identify the optimal combination of matching approach and gene sets, we implemented a three-step validation strategy (**Figure 1**): first, matching subsets of cells of identical type within the same scRNA-seq dataset, second, comparing cell types across independent scRNA-seq references while retaining the most accurate combinations, and third, testing on microdissected kidney tissue (positive controls) and non-kidney samples (negative controls). The most accurate approach was then applied to assess commonly used kidney tubular cell culture models for their similarity to kidney tubular cell types. The complete methodology is detailed in the **Supplementary Methods**.

**Figure 1:**
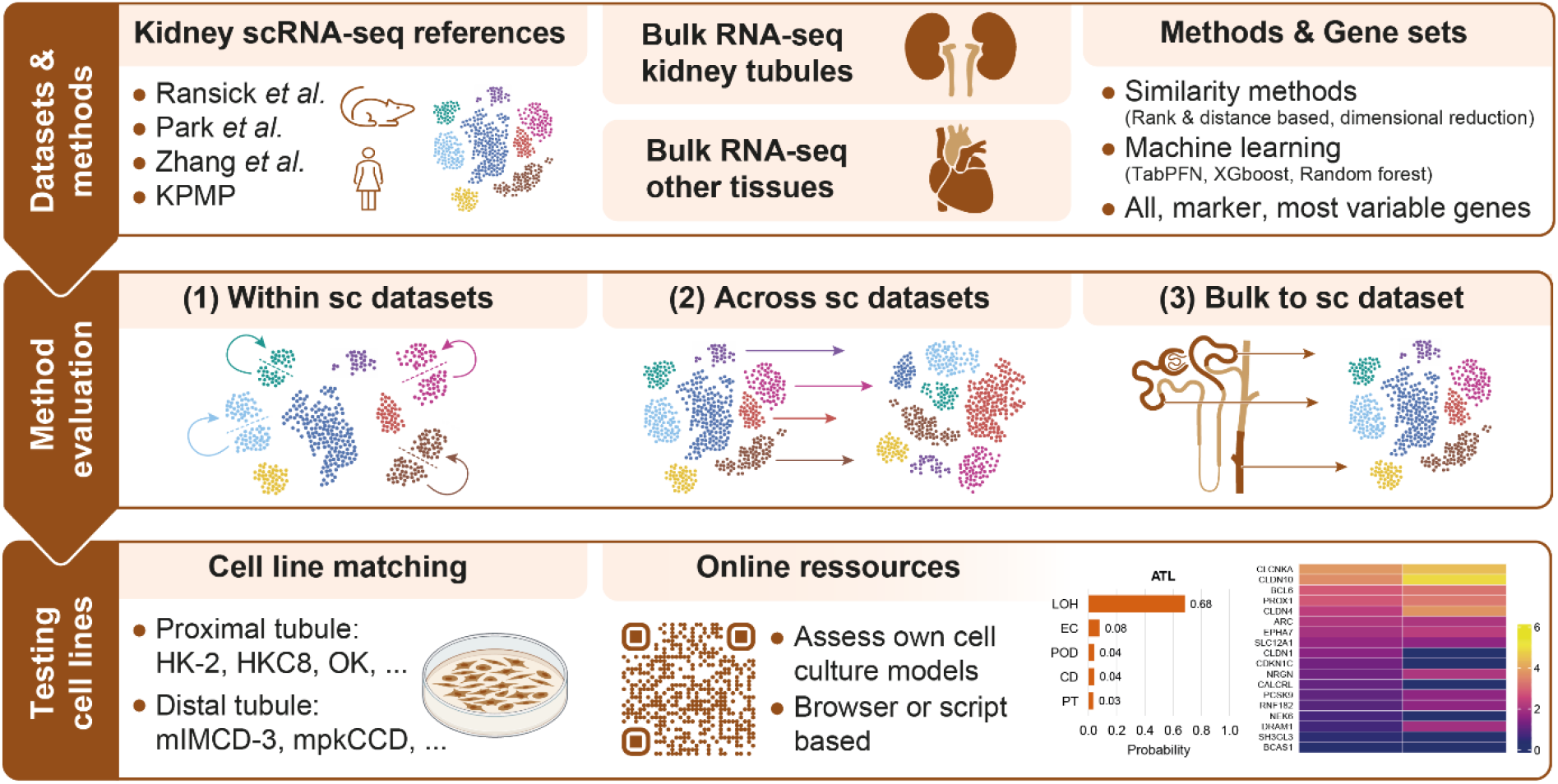
Study workflow. Reference transcriptomic profiles for kidney cell types were derived from four human and mouse scRNA-seq datasets. Bulk RNA-seq data from kidney and other tissues were matched to these references using different gene sets (global expression and marker gene lists) and various similarity and machine learning approaches. The optimal combination of gene set and matching approach was determined by sequentially matching pseudobulk samples derived from cell types within a given scRNA-seq dataset (1), across scRNA-seq datasets (2), and by matching bulk RNA-seq data from primary kidney and other tissues to the scRNA-seq references (3). The best-performing approaches were used to characterize commonly used cell culture models and are provided online for the scientific community (link will be added after peer reviewed publication).

## Results

### Matching cell types within and across scRNA-seq reference datasets

We randomly split cells from each cell type within each reference dataset into two groups, generated aggregated expression profiles for each, and assessed how well “group 1” profiles were correctly matched to their corresponding “group 2” cell types. While all ML classifiers performed well, TabPFN and Random Forest achieved over 99% accuracy regardless of the gene set used. Over 97% accuracy was achieved with all three similarity methods in combination with the curated kidney marker gene lists (**Figure 2a, Supplementary Table 4**). Next, cell types were matched across different scRNA-seq reference datasets, causing median accuracy to decline to 56% from 92% observed in matches within-reference (**Figure 2b**). Again, Random Forest, TabPFN, and the three similarity methods performed best, particularly in combination with kidney marker genes. Cross-dataset alignment differed substantially between the references: KPMP showed the poorest alignment with other scRNA-seq references across all method-gene set combinations (median accuracy 49%), while Ransick *et al*. performed best (median accuracy 59%, **Supplementary Table 5**). Overall, matching within and across scRNA-seq references demonstrated that all three similarity measures and the ML approaches performed best when applied with either all detected genes or the marker gene sets. Cell type matching based on most variable genes or by combining similarity methods with prior dimensional reduction performed worse and was excluded from subsequent analyses.

**Figure 2:**
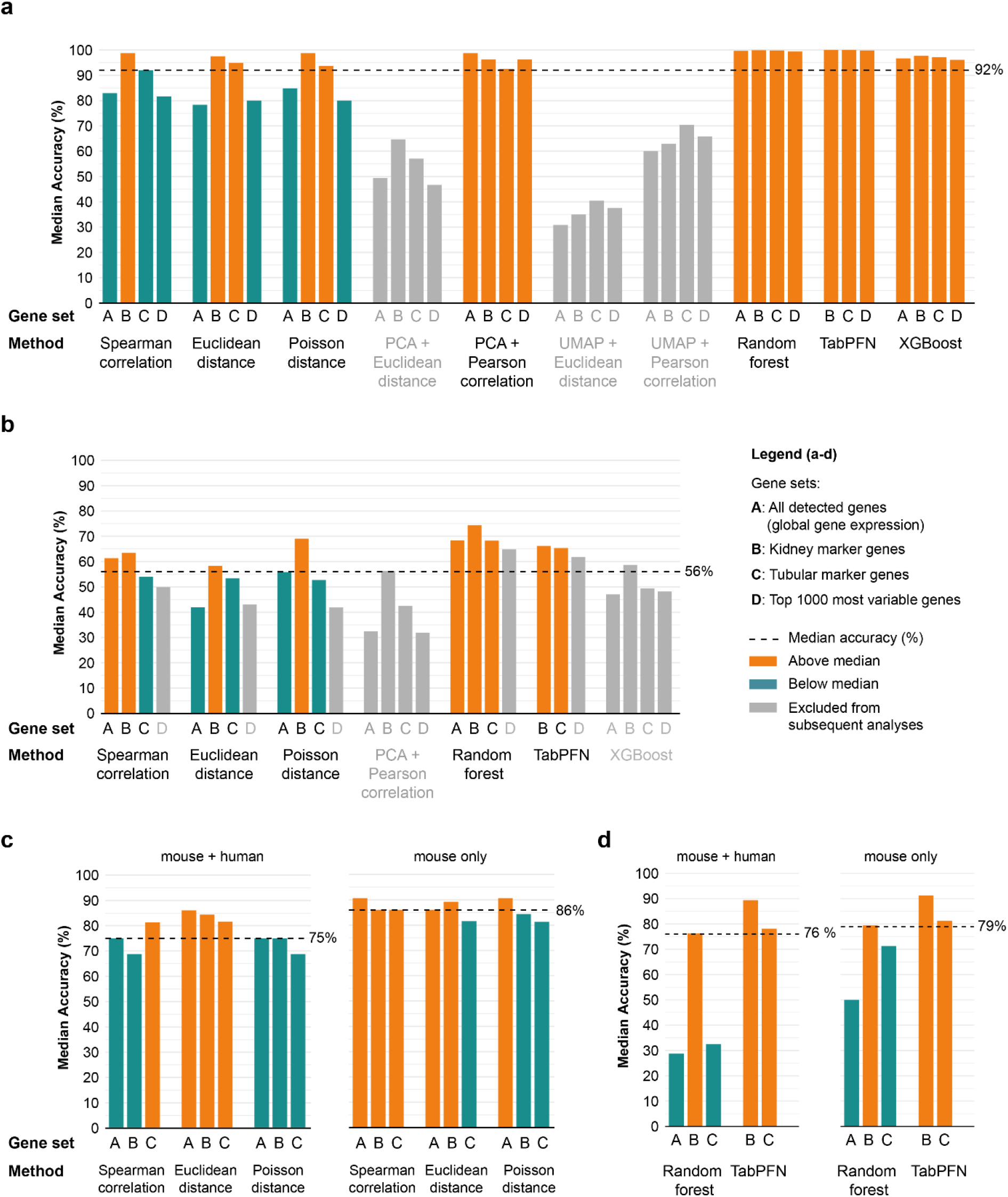
Evaluation of matching methods. Panels show median matching accuracies (%) for tested method-gene set combinations across all included kidney scRNA-seq reference datasets. TabPFN was not tested with all detected genes (A) as this exceeds the model’s feature limit. **(a)** Within-dataset matching accuracy (matching subsets of identical cell clusters from the same scRNA-seq reference). **(b)** Between-reference matching accuracy (matching corresponding clusters across different scRNA-seq references). **(c)** Matching accuracy of 16 microdissected mouse bulk RNA-seq tissue samples to all references using similarity methods only (left: all scRNA-seq references; right: mouse references only). **(d)** Matching accuracy of 16 mouse bulk RNA-seq tissue samples to all references using machine learning methods only (left: all scRNA-seq references; right: mouse references only). Abbreviations: PCA, principal component analysis; TabPFN, Tabular Prior-data Fitted Network; UMAP, Uniform Manifold Approximation and Projection.

### Matching of microdissected primary kidney tissue samples

We further assessed the matching accuracy of the 15 remaining method-geneset combinations based on primary kidney tissue using bulk RNA-seq data from 14 microdissected mouse tubule samples^25^ as well as mouse kidney medulla and cortex tissue. Median accuracy across combinations with similarity methods was 75% and further increased to 86% when only murine single-cell references were used (**Figure 2c, Supplementary Table 6**). Accuracy was higher when broader gene sets (all genes and mg_all) were used. ML methods, particularly TabPFN, maintained stable accuracy irrespective of species-matched references (**Figure 2d**). Overall, near-perfect matching accuracy was achieved for most tubular segments across methods and gene sets, except descending thin limb (DTL1–3), assigned to varying cell types depending on the sc reference, and connecting tubule (CNT) sometimes matched to reference cell types of the collecting duct (**Supplementary Tables 7 and 8)**. To illustrate these findings, **Figure 3a** shows the successful TabPFN-based matching using mg_all of bulk RNA-seq data from microdissected proximal tubule segment 2 (PTS2), ascending thin limb (ATL), and cortical collecting duct (CCD) to their corresponding pooled reference cell type. **Figure 3b** illustrates that these results align well with the correlation of expression-based gene ranks for the marker genes included in this analysis.

**Figure 3:**
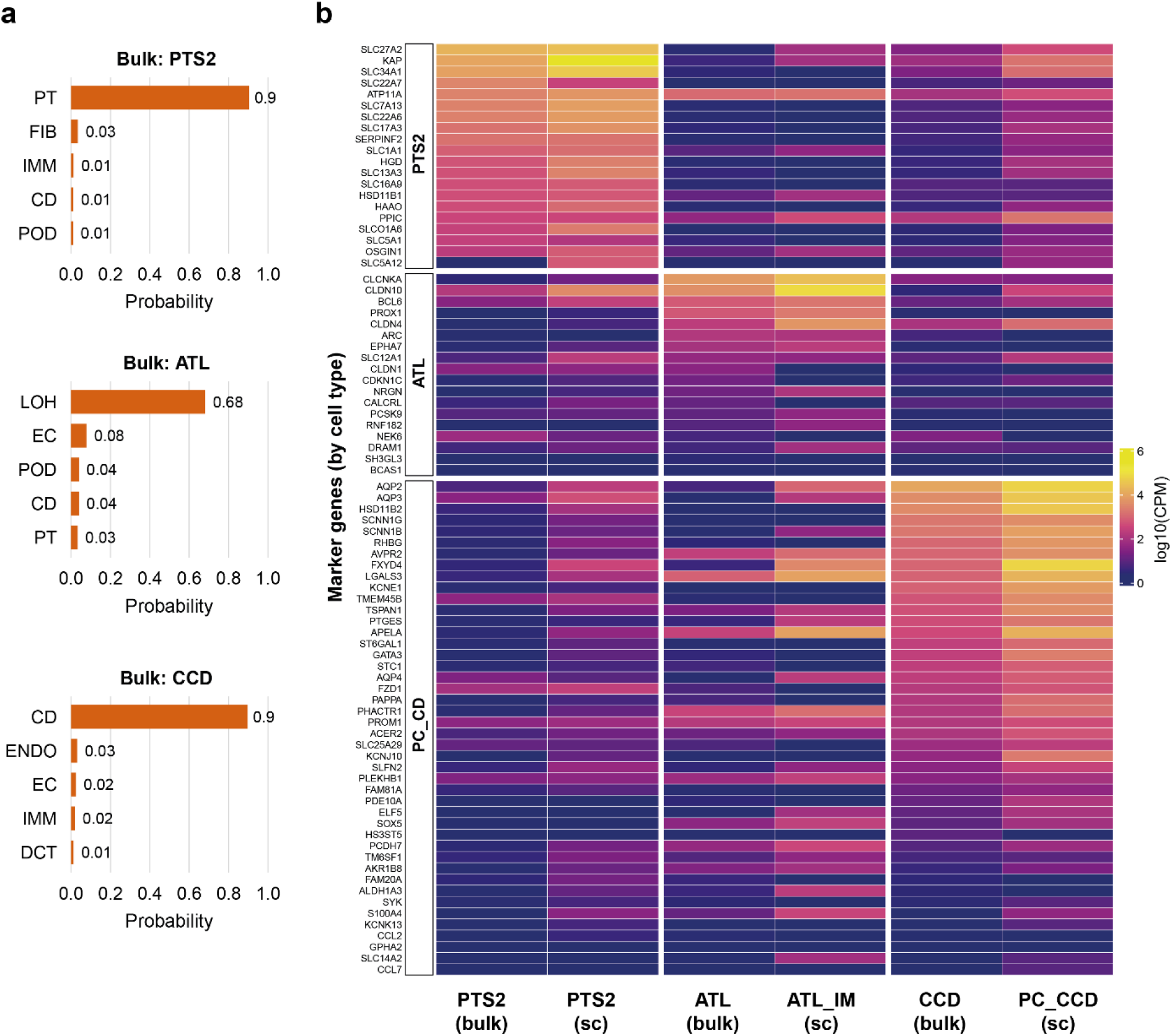
Matching results for microdissected tubular segments. **(a)** TabPFN-based matching results for bulk RNA-seq data from three microdissected kidney tubule segments (PTS2, ATL, CCD) using our kidney marker gene set (mg_all). The top five reference dataset-derived cell types are shown based on TabPFN probability output. **(b)** Expression of marker genes from mg_all for PTS2, ATL, and CCD as well as collecting duct principal cells (PC_CCD) in the bulk RNA-seq samples (bulk) and in the scRNA-seq reference (sc) from Ransick *et al*. (pseudobulk). Marker genes for each cell type are ordered by expression magnitude (log10(CPM)) in the bulk data. Gene expression ranks show good concordance between bulk and scRNA-seq data for matching cell types. Abbreviations: ATL, ascending thin limb; CCD, cortical collecting duct; CPM, counts per million; PTS2, proximal tubule segment 2.

To assess specificity, bulk RNA-seq data from non-kidney tissues (non-kidney fibroblasts, heart, brain, blood monocytes) were matched as negative controls. Spearman correlation showed the lowest false-positive rate for tubular cell types among similarity methods across gene sets (10%, **Supplementary Table 9**), while both ML approaches, TabPFN and Random Forest, achieved an even lower rate of 0%.

Overall, although all three similarity approaches display similarly good performance, Spearman correlation-based cell type matching, particularly in combination with the broad kidney marker gene list showed the most accurate and specific results. Among the ML algorithms, TabPFN shows the best combination of accuracy and specificity when used with the respective sets of marker genes and additionally quantifies the uncertainty in the prediction.

### Cell Line Matching

Finally, we assessed the transcriptomic similarity of various kidney cell lines with different culture conditions and treatments to cell types in our scRNA-seq references using TabPFN and rank-based Spearman correlation which is summarized in **Table 1** from the detailed results in **Supplementary Tables 10 and 11**. Focusing on kidney marker gene-based comparisons, the commonly used proximal tubule OK and HK-2 cell lines were most similar to proximal tubule cells. Similarity further increased when OK cells were exposed to shear stress in culture, mimicking the fluid flow present in their native environment. Other proximal tubule cell lines (HKC-11, HKC-8, HPTC) showed inconsistent matching results across methods and gene sets, suggesting variable preservation of proximal tubule identity. Similarly, HUPEC cells derived from podocytes did not consistently match any specific reference cell type, including podocytes. Among more distal nephron cell lines, C125 derived from kidney medulla matched loop of Henle and collecting duct cells correctly and consistently, confirming previous findings^26^. Cell lines derived from the cortical collecting duct (mpkCCD) or inner medullary collecting duct (mIMCD3) consistently matched loop of Henle by TabPFN and collecting duct by Spearman correlation, regardless of two- or three-dimensional culture conditions or *Pkd1* knockout (**Table 1**). Primary cells of the inner medullary collecting duct (IMCD) showed the same pattern. Notably, increased medium osmolality shifted TabPFN probabilities toward loop of Henle for both mpkCCD (300 vs. 600 mOsm) and primary IMCD cells (300 vs. 900 mOsm), suggesting that osmolality-induced broader transcriptomic changes are captured by this approach.

**Table 1:** Matching results for tubular cell lines. Matching results for various cell lines derived from proximal tubule, kidney medulla, and collecting duct across different species under the indicated treatment and culture conditions. The TabPFN column shows the most probable cell type match from a model trained on all four scRNA-seq reference datasets. The Spearman correlation column shows the most similar cell type based on correlation-based matching against all four scRNA-seq references; if matching was consistent across two or more references, the respective cell type is shown, otherwise “undecided” is displayed. Both TabPFN and Spearman correlation of gene ranks were applied using the kidney marker gene list (mg_all; see Methods). Abbreviations: api, apical; baso, basolateral (cells cultured on transwell filters); CD, collecting duct; IMM, immune cells (see Supplementary Table 13 for details); KO, knockout; LOH, loop of Henle; PT, proximal tubule; WT, wild type.

| Cell line | Celltype or tissue of origin | Species | Treatment | TabPFN (probability) | Spearman correlation |
| --- | --- | --- | --- | --- | --- |
| OK | Proximal tubule | opossum | WT | IMM (0.35) | PT |
|  |  |  | Shear stress | PT (0.31) | PT |
| HK-2 | Proximal tubule | human | WT | IMM (0.27) | PT |
| HKC-11 | Proximal tubule | human | WT | IMM (0.44) | undecided |
| HKC-8 | Proximal tubule | human | WT | IMM (0.38) | undecided |
| HPTC.05.LTR | Proximal tubule | human | WT | IMM (0.34) | undecided |
| HUPEC_1 | Podocytes | human | WT | PT (0.25) | undecided |
| HUPEC_2 | Podocytes | human | WT | PT (0.29) | undecided |
| C125 | Kidney medulla | mouse | *Pkd2* KO, passage 4 | LOH (0.32) | CD |
|  |  |  | *Pkd2* KO, passage 7 | LOH (0.28) | CD |
| mpkCCD | Cortical collecting duct | mouse | 300mOsm (api & baso) | LOH (0.22) | CD |
|  |  | mouse | 600mOsm (api & baso) | LOH (0.29) | CD |
|  |  | mouse | 300mOsm (api) / 600mOsm (baso) | LOH (0.25) | CD |
|  |  | mouse | 600mOsm (api) / 300mOsm (baso) | LOH (0.31) | CD |
| mIMCD-3 | Inner medullary collecting duct | mouse | *Pkd1* KO 1 | LOH (0.50) | CD |
|  |  |  | *Pkd1* KO 2 | LOH (0.53) | CD |
|  |  |  | WT | LOH (0.46) | undecided |
|  |  |  | WT 3D culture | LOH (0.48) | CD |
| IMCD (primary cells) | Inner medullary collecting duct | rat (f) | 300mOsm | LOH (0.44) | CD |
|  |  | rat (m) | 300mOsm | LOH (0.45) | CD |
|  |  | rat (f) | 900mOsm | LOH (0.56) | CD |
|  |  | rat (m) | 900mOsm | LOH (0.55) | CD |

## Discussion

We developed a transcriptome-based approach to match kidney cell lines and tissues to scRNA-seq reference cell types – an application for which, as described in the introduction, no dedicated approach has previously been established. Gene expression rank-based Spearman correlation and TabPFN, a machine learning foundation model, emerged as the most accurate and specific methods, particularly when combined with curated kidney marker gene lists. Both methods achieved matching accuracies of up to 90% for microdissected primary murine kidney tissue when species-matched references and kidney-specific marker genes were used (**Figure 2c-d**), demonstrating reliable cell type identification from bulk RNA-seq data.

Considerable variance in matching accuracy was observed across the four tested scRNA-seq references, a finding consistent with well-documented heterogeneity in cell type annotations, experimental protocols, and biological variation between datasets^27,28^. This inter-reference variability is reflected in the decline in cross-reference matching accuracy (56% vs. 92% within-reference, **Figure 2a-b**) and underscores the importance of evaluating multiple references rather than relying on a single dataset. Among proximal tubule cell lines, matching results were heterogeneous. OK cells showed the strongest similarity to proximal tubule, consistent with previous findings that OK cells retain more proximal tubule characteristics than other commonly used lines^8^. Notably, similarity increased when OK cells were cultured under fluid shear stress^29^, which induces differentiation toward a more native-like phenotype. Other proximal tubule lines (HKC-11, HKC-8, HPTC) and podocyte-derived HUPEC cells showed inconsistent matching, suggesting de-differentiation in culture. Distal nephron and collecting duct cell lines showed more consistent results with Spearman correlation-based matching. Interestingly, TabPFN assigned collecting duct-derived cell lines (mpkCCD, mIMCD-3, primary IMCD) to loop of Henle rather than collecting duct, with increasing probability at higher medium osmolality. This pattern may reflect corticomedullary gene expression gradients previously described in single-cell datasets^30^ and bulk datasets^31^: the loop of Henle cell type learned by TabPFN likely incorporates transcriptomic features of the deep medullary, high-osmolality environment, whereas collecting duct reference profiles may be dominated by more cortical collecting duct cells. This interpretation is supported by the osmolality-dependent shift in matching probabilities and highlights how culture conditions that mimic the native microenvironment are captured by our approach.

Key strengths of this study include our rigorous three-step validation strategy, the evaluation of both human and mouse references demonstrating improved accuracy with species-matching, and the provision of curated marker gene lists that outperformed highly variable genes. Importantly, our approach captures broader transcriptomic shifts from physiological challenges (osmolality, shear stress) rather than single-gene perturbations, complementing traditional marker-based assessments.

Regarding limitations, matching accuracy varied considerably across scRNA-seq references, which can yield inconsistent results for cell types that are sparsely represented or have overlapping transcriptomic signatures in available references – notably the descending thin limb and connecting tubule, whose distinct identities are difficult to resolve when reference coverage is limited. Additionally, our approach is inherently constrained to cell types present in the reference datasets used. TabPFN partially addresses this by training on pooled cell types across multiple references, improving robustness to individual reference biases, though inter-reference variability remains an intrinsic challenge that cannot be fully resolved.

The two recommended approaches offer complementary advantages: Spearman correlation is transparent, fast, and enables comparison against individual references, while TabPFN leverages pooled training and provides probability scores with greater discriminative spread. We provide an accessible web-based tool for routine use (link will be added after peer reviewed publication) and comprehensive scripts for TabPFN-based matching (link will be added after peer reviewed publication). Together, these resources enable the kidney research community to make informed decisions about cell culture model selection, stability, and interpretation. Future extensions could include application to kidney organoids and iPSC-derived models, as well as integration of additional scRNA-seq references as they become available.

## Acknowledgements

The work of M.S., M.K., L.W., Y.L., G.G., P.S., S.H., O.B., and A.K. was funded by German Research Foundation (DFG) project ID 431984000 (SFB 1453). Germany’s Excellence Strategy (CIBSS, EXC-2189, project ID 390939984) supported the work of M.S., P.S., M.K. and A.K.

The work of P.S. was supported by the DFG project-ID 530592017 (SCHL 2292/3–1). The work of S.B. and P.S. was supported by the DFG project-ID 499552394 – CRC 1597 Small Data.

The creation of the C125 cell line and the work of O.M.W and H.J.J was funded by NIH U54 1U54DK126114.

The work of H.J.J was supported by the Korea Health Technology R&D Project through the Korea Health Industry Development Institute (KHIDI), funded by the Ministry of Health & Welfare, Republic of Korea (RS-2025-25410994), and the Edward S. Kraus Scholar Award, Johns Hopkins University School of Medicine Division of Nephrology.

## Data Sharing Statement

Comprehensive summary data are provided in the supplementary materials, including detailed information on the included scRNA-seq and bulk RNA-seq datasets and cell types (**Supplementary Tables 1**,**2, 12 and 13**), curated marker gene lists (**Supplementary Table 3**), and detailed matching results for all analyses described in the results section (**Supplementary Tables 4-11**). Code used for machine learning-based matching (TabPFN) is available in a GitHub repository accompanying this publication (link will be added after peer reviewed publication). Here we also present a lightweight implementation of Spearman correlation-based cell type matching using our scRNA-seq references and marker gene lists as a ShinyLive app that can be executed locally in any web browser with user-provided datasets. The app is directly accessible at link will be added after peer reviewed publication. Previously published scRNA-seq and bulk RNA-seq datasets used in this study are available from their original sources as referenced in the Supplementary Tables. Additional data are available from the corresponding author upon reasonable request.

## Author Contributions

Design and conceptualization of this study: M.K., P.S., A.K., S.H.

Transcriptome data of cell lines: O.M.W., H.J.J., B.E., L.W., M.K.

Bioinformatics and statistical analysis: M.S., Y.L., O.B., S.H.

Interpretation of Results: M.S., S.B., P.S., A.K., S.H.

Wrote the manuscript: M.S., S.B., S.H.

Critically read and approved the manuscript: M.S., S.B., O.B., Y.L., G.G., B.E., O.M.W., H.J.J., M.K., L.W. F.H., P.S., A.K., S.H.

## Generative AI language models

During the preparation of this work, the authors used generative AI language models (Claude Sonnet 4.5 and Opus 4.6, Anthropic) to improve the readability and language of the manuscript. After using this tool, the authors reviewed and edited the content as needed and take full responsibility for the content of the publication.

